# Assembly mechanism of the AIM2 inflammasome sensor revealed by single-molecule analysis

**DOI:** 10.1101/2022.09.21.508942

**Authors:** Meenakshi Sharma, Eva de Alba

## Abstract

Pathogenic dsDNA prompts AIM2 (Absent In Melanoma 2) assembly leading to the formation of the inflammasome, a multimeric complex that triggers the inflammatory response. The recognition of foreign dsDNA involves AIM2 self-assembly concomitant with dsDNA binding. However, we lack mechanistic and kinetic information on the formation and propagation of the assembly, which can shed light on innate immunity’s time response and specificity. Using correlative optical traps and fluorescence microscopy, we determine here the association and dissociation rates of the AIM2-DNA complex at the single-molecule level. We identify distinct mechanisms for oligomer growth via the binding of incoming AIM2 molecules to adjacent dsDNA or direct interaction with bound AIM2 assemblies, thus resembling primary and secondary nucleation processes. Through these mechanisms, AIM2 oligomers increase at least fourfold their size in seconds. Finally, our data indicate that single AIM2 molecules do not diffuse/scan along the DNA, suggesting that oligomerization depends on stochastic encounters with DNA and/or DNA-bound AIM2 molecules.

---

The innate immune system recognizes cues associated with cellular damage and molecular patterns arising from invading pathogens^1^. Recognition by sensor proteins triggers the assembly of large signaling platforms known as inflammasomes that activate inflammatory caspases^2–4^. Inflammasome formation involves sensor oligomerization^5–7,^ leading to the self-assembly of the adaptor protein ASC into the so-called “ASC speck”^8,9^. ASC recruits the effector caspase 1^2,10^, increasing its local concentration and promoting caspase activation^11^, thus resulting in the maturation of proinflammatory cytokines^12^ and cell death by pyroptosis and PANoptosis ^13,14^. In the process of inflammasome activation, a filamentous punctum (ASC speck) with a diameter of ∼ 0.5 - 1 µm forms via self-association and oligomerization of multiple protein components (**Figure 1a**)^9^. At the molecular level, it has been shown that the inflammasome adaptor ASC and its isoform ASCb^8^ with two oligomerization Death Domains, PYD and CARD, can polymerize into different macrostructures^15–18^. Via homotypic interactions, ASC connects PYD-containing sensors^19^ and procaspase 1 to facilitate speck assembly and activation^20^.

**Figure 1.**
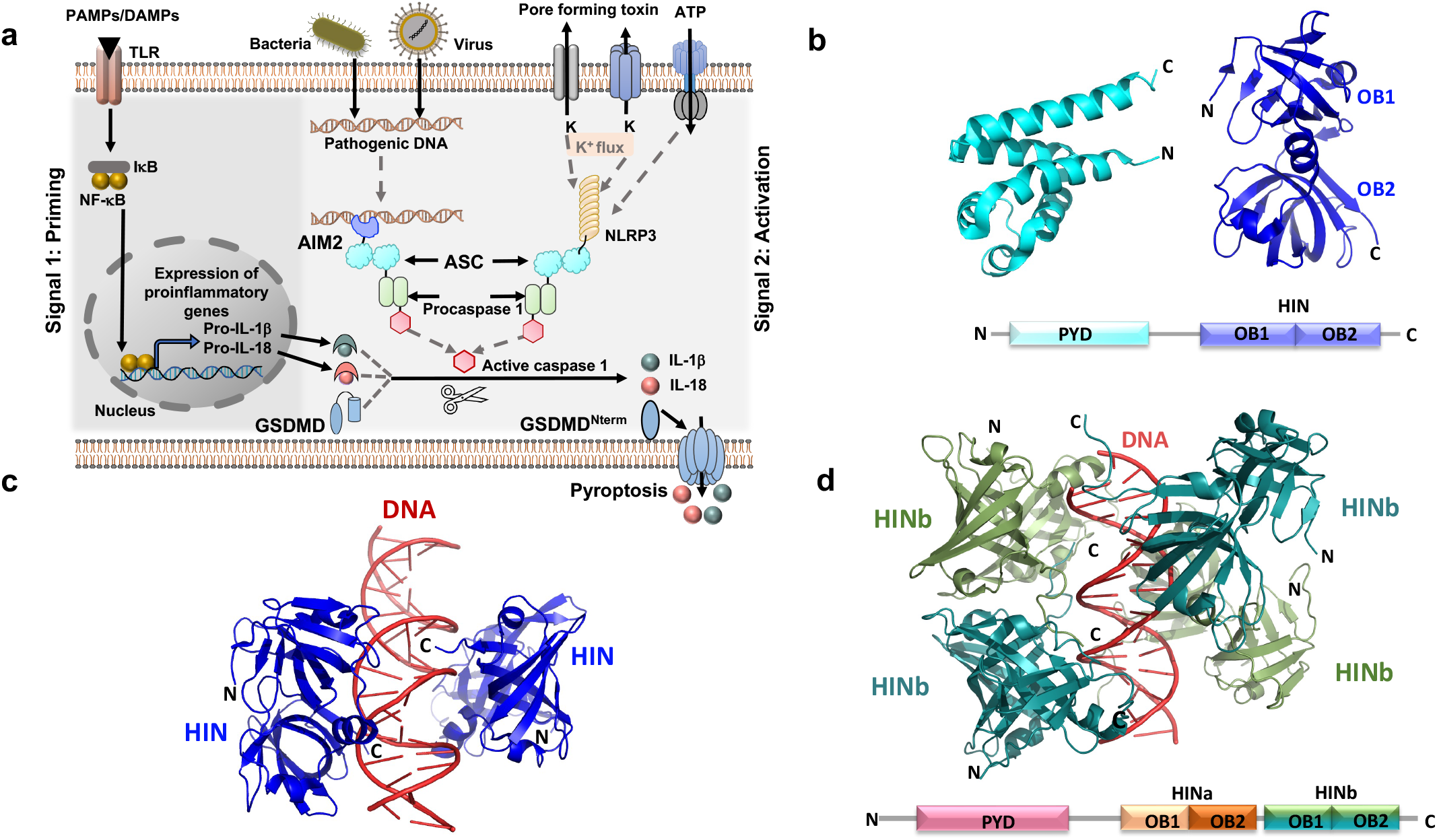
Inflammasome operating mode and structure of dsDNA inflammasome sensors. **a** Priming and activation of NLRP3 and AIM2 inflammasomes. The adaptor ASC and maturation products from procaspase 1 activation are indicated. **b** Domain organization and individual three-dimensional structures of AIM2^HIN 25^ and AIM2^PYD 48,49^. **c** Three-dimensional structure of the complex between dsDNA and AIM2^HIN 25^. **d** Three-dimensional structure of the complex between dsDNA and the HINb domain of IFI16^25^. IFI16 domain organization (bottom).

Inflammasome sensors show specificity for different molecular patterns. For instance, foreign dsDNA activates the sensors AIM2 and IFI16^21–24^. Both sensors carry an N-terminal PYD for self-assembly and polymerization with ASC (**Figure 1b, d**), leading to the formation of the inflammasome, and C-terminal HIN (Hematopoietic, Interferon-inducible, Nuclear localization) domain(s) for DNA binding. However, AIM2 is a cytosolic sensor, whereas IFI16 is the only sensor identified thus far that recognizes foreign DNA in the nucleus^25,26^. Detailed functional and structural studies of complexes between dsDNA and the DNA binding domains of the cytosolic and nuclear sensors explain the lack of sequence specificity, as the intermolecular interactions involve the dsDNA phosphate backbone (**Figure 1c, d**)^25^. Importantly, the authors indicate that dsDNA serves as an oligomerization platform for the inflammasome and inform on the estimated size of the oligomers. These studies are complemented by functional cell assays showing that optimal IL-1β induction is achieved when transfecting ∼ 80 bp dsDNA, which will be able to host up to 20 HIN domains based on the crystallographic structures^25^ (**Figure 1c, d**).

Equally elegant studies on the function and operating modes of AIM2 and IFI16 found that DNA binding and sensor self-association are integrated and cooperative processes^26,27^. The PYD domain was found to be essential for these functions, and specifically required for strong binding to dsDNA and polymerization in the presence of excess dsDNA. These studies show that AIM2-DNA and IFI16-DNA binding affinity depends on the DNA length, as the affinity increases steeply for dsDNA longer than a threshold of ∼ 70-bp (hosting ∼ 6 AIM2 protomers) until reaching a maximum value for ∼ 280 bp-DNA (hosting ∼ 24 AIM2 protomers)^27^. This work thus indicates that the DNA acts as a molecular ruler for AIM2 inflammasome assembly following a switch-like mechanism ^27^.

Furthermore, the interaction between the nuclear sensor IFI16 and dsDNA has been studied using single-molecule fluorescence imaging by TIRF microscopy (Total Internal Reflection Fluorescence)^28^. This study shows single IFI16 molecules diffusing several µm along the λ-phage dsDNA. IFI16 scans the dsDNA to find other molecules already bound to DNA for oligomerization. A sufficiently long stretch of free dsDNA is required for scanning and oligomerization, thus elegantly explaining how IFI16 discriminates between self- and foreign-DNA, as the former does not expose sufficiently long dsDNA fragments available for self-assembly due to nucleosome packing^28^.

Despite the challenges associated with protein oligomerization, combined efforts using a variety of biophysical, biochemical and microscopy techniques are significantly advancing our understanding on AIM2 inflammasome formation^25,27–30^. However, there are unresolved questions on mechanistic and kinetic aspects of the AIM2-DNA assembly process. First, we do not know whether AIM2 forms individual, distinct oligomers on the DNA, and in this case, whether oligomers of different sizes can coexist and what their shapes are. Second, what are the association and dissociation rates of a single AIM2-DNA complex? Third, how fast do oligomers grow and what mechanisms are followed for oligomerization and propagation? And finally, since AIM2 does not need to discriminate between self- and foreign-DNA, does the cytosolic sensor follow the same scanning mechanism as the nuclear sensor IFI16? Here, we provide answers to these questions using correlative optical traps and confocal fluorescence microscopy to study the initial stages of the AIM2 inflammasome.

## Results

### AIM2 forms distinct oligomers of different sizes and shapes bound to a single dsDNA molecule

The typical experimental setup using correlative optical traps and confocal fluorescence microscopy is shown in **Figure 2a**. A single λ-phage dsDNA molecule (∼ 16.5 µm long) is tethered between two optically trapped beads (Ø ∼ 3 µm) and moved to the protein channel. Two-dimensional (2D) images are acquired by laser scanning confocal fluorescence microscopy. In seconds, fluorescent AIM2 clusters of various sizes populate multiple positions of the dsDNA molecule at sub-nanomolar to low nanomolar protein concentrations (∼ 0.2 nM – 14 nM) under physiological salt concentration (160 mM KCl) (**Figure 2b**). In some instances, small AIM2 clusters span the DNA molecule and in other cases, only a few large clusters appear. Large and small clusters are observed together (**Figure 2b**). The multiple binding events along the dsDNA molecule suggest that AIM2 lacks sequence specificity, as demonstrated by the X-ray structural studies on the AIM2^HIN^-dsDNA complex^25^.

**Figure 2.**
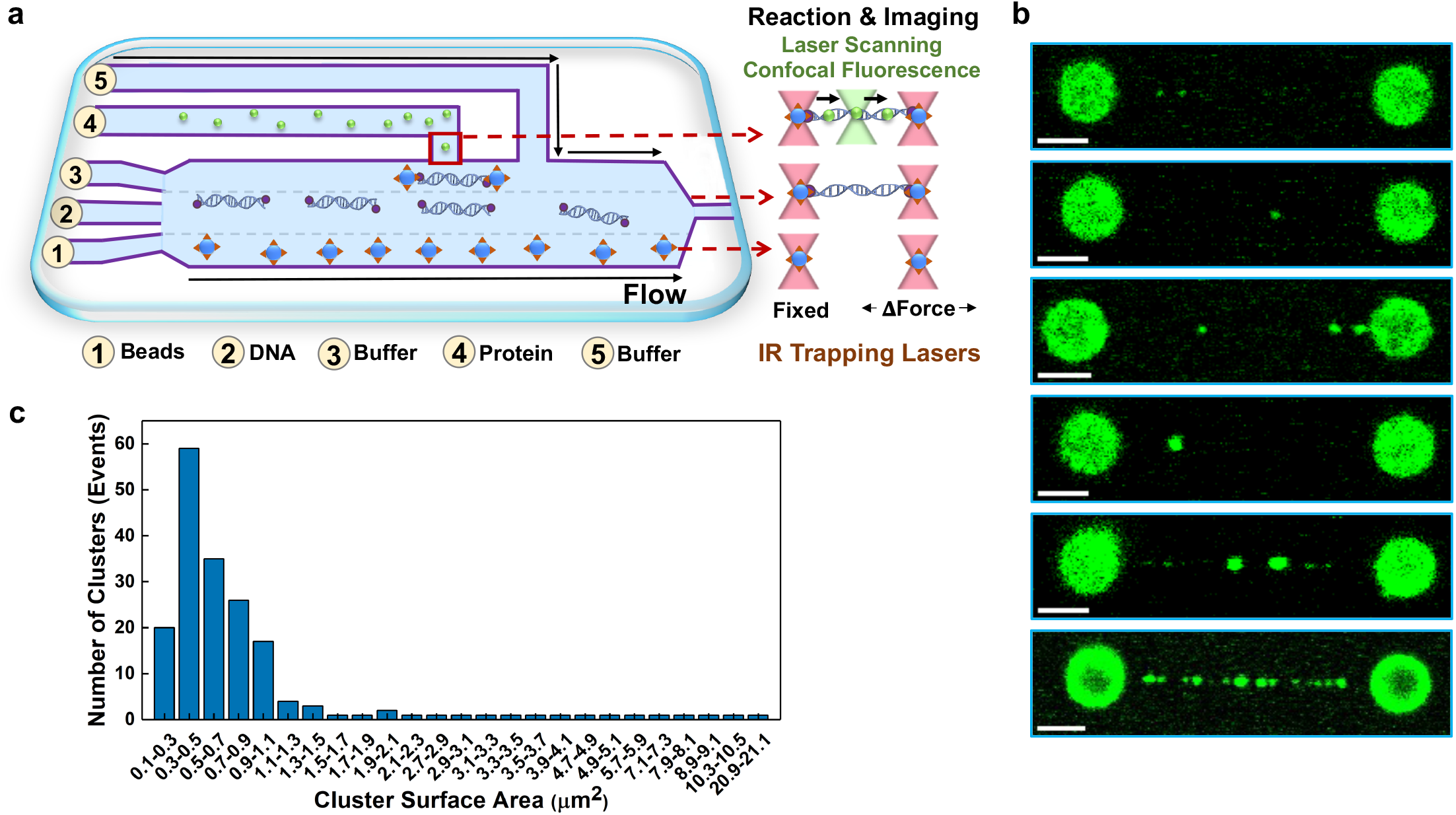
AIM2 forms distinct oligomers of different sizes and shapes bound to a single dsDNA molecule. **a** Schematic representation of a typical experimental setup, showing the microfluidics system in the flow cell: streptavidin coated beads in channel 1, biotinylated phage-λ dsDNA in channel 2 and fluorescent AIM2 in channel 4. **b** Representative examples of two-dimensional scans obtained seconds after exposing the dsDNA to the protein channel. **c** Cluster size distribution based on surface area (N= 183).

An analysis of the surface area distribution of single AIM2 particles reveals that the clusters typically fall in the 0.1 − 1.1 µm^2^ range (**Figure 2c**). The smallest cluster size observed (0.1 µm^2^) is diffraction limited (∼ 175 nm at λ = 525 nm) and clusters in the 0.3 – 0.5 µm^2^ interval are the most abundant. Clusters larger than 0.9 – 1.1 µm^2^ are increasingly infrequent (**Figure 2c**). It must be noted that oligomer surface area is only a partial characteristic obtained from the analysis of 2D images of 3D particles. The confocal images also show that AIM2 clusters adopt different shapes. Small clusters are typically round, and larger clusters are often asymmetric, suggesting that cluster growth may happen by AIM2 binding directly to an attached oligomer without interacting with the DNA. Overall, this analysis shows that AIM2 oligomers of different sizes and shapes coexist upon DNA binding.

### AIM2 oligomers attached to the single dsDNA are typically smaller than 25 molecules

We have estimated the number of molecules in the different clusters using fluorescence intensity relative to the intensity produced by a single fluorophore. Several assumptions were made to correlate fluorescence intensity with the number of fluorophores. Specifically, we assume that the detector response is close to linear due to the low dead time (35 ns) of the Avalanche Photodiode Detector (APD). This dead time results in ∼ 6% underestimation of photon counts for a cluster of 10 emitting fluorophores, leading to an error of ∼ 1 fluorophore in the 10-fluorophore cluster. An approximately linear APD response is achieved by working under conditions that avoid detector saturation (i.e., low laser power (10%) and low number of photons detected due to the confocal setup). In addition, both the numerical aperture of the objective and the confocal microscopy setup restrict the angles at which photons are detected. This effect may be ignored for fluorophores with isotropic rotation. However, isotropic motion might be compromised in the presence of AIM2 oligomerization, thus photons emitted by fluorophores positioned at the appropriate angles might have greater chances of being detected. Our estimations do not consider this effect. Additionally, we can safely assume that laser excitation (at constant power) is uniform across the ROI (region of interest) by scanning the confocal plane. With respect to the optical axis, we assume the detector collects photons emitted by assembled molecules laying within the Z-axis resolution of the microscope (∼ 1µm). This assumption might result in a slight underestimation of the number of molecules even if clusters are rarely larger than 1 µm in diameter due to variations in excitation along the Z-axis.

In addition, correction factors need to be applied to consider intensity decay due to photodepletion and fluorophore labeling efficiency. Evidence show that fluorophore clustering affects the emission properties of individual fluorophores^31,32^. Therefore, the total emitted light of fluorophores clustered in proximity is frequently enhanced or decreased relative to the expected intensity from the total number of fluorophores^31,33^. For these reasons, fluorophore clustering poses challenges in determining the stoichiometry of protein complexes based on fluorescence intensity^33–36^. For AIM2, the fluorescence intensity emitted by protein oligomers bound to dsDNA decreases with time due to photodepletion processes (**Supplementary Figure 1a**). Decay rates of the fluorescence intensity produced by AIM2 clusters vary from 0.2 s^-1^ to 0.5 s^-1^ with an average of 0.43 s^-1^ for clusters emitting less than 1,000 photon counts. Based on these decay rates, the effect of photodepletion is negligible during the short excitation time of the scanning laser (∼ 5 ms for an ROI of 1µm^2^). In addition, fluorescence intensity was corrected to consider a labeling efficiency of ∼ 80% (Methods).

After this correction, the initial fluorescence intensity divided by the intensity emitted by a single fluorophore allows to estimate the number of molecules per oligomer. The photon counts emitted per fluorophore were determined by the identification of photodepletion steps in fluorescence intensity decays (Methods, **Supplementary Figure 1a**)^37,38^. The resulting histogram distribution shows that 11 photon counts are more frequent, thus corresponding to a single fluorophore (**Supplementary Figure 1b**). As expected, the intensity of single fluorophores is mainly steady with time (**Supplementary Figure 1c, d**).

The distribution of the number of molecules per oligomer resulting from the analysis of 183 clusters shows that oligomers smaller than 25 molecules are more abundant (**Figure 3a**). There is no clear predominance of a specific oligomer size within this range, except for clusters composed of 13-15 molecules being slightly more frequent (**Figure 3b**). Within associated errors, these results are in accord with previous studies indicating a preferred oligomer size of 20-24 AIM2 molecules^25,27^. The direct visualization of individual oligomers reveals the coexistence of small and large AIM2 clusters (**Figure 2b, c & Figure 3**). Thus, our results provide additional insight into the all-or-none process suggested for AIM2-DNA binding based on the sigmoidal response of AIM2 affinity for different lengths of dsDNA^27^.

**Figure 3.**
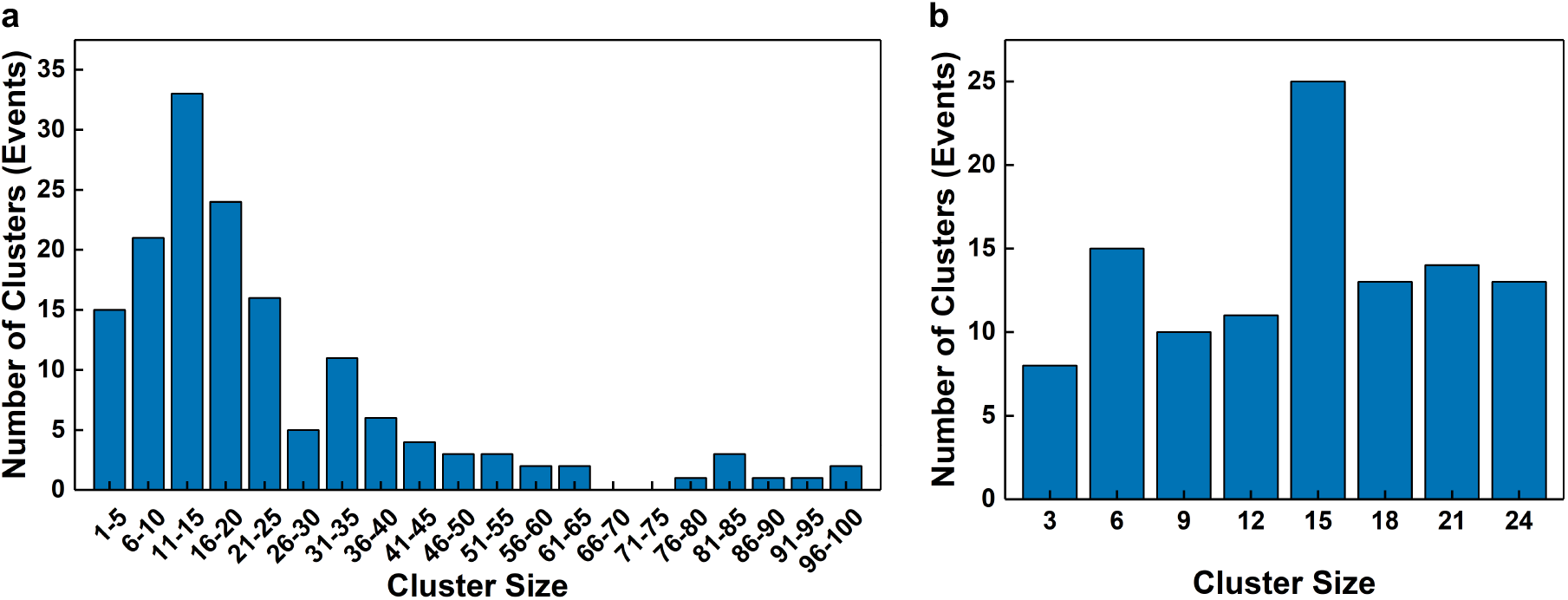
AIM2 oligomers attached to a single dsDNA are predominantly smaller than 25 molecules. **a** Cluster size distribution based on the number of molecules up to 100 molecules and **b** up to 25 molecules (N = 183). Oligomers of 13-15 molecules are slightly more frequent.

Clusters larger than 50 AIM2 molecules are less frequent (**Figure 3a**). Large clusters composed of ∼ 1,000 to ∼ 5,000 molecules have been observed both bound to dsDNA and trapped together with the beads when there is no DNA (**Supplementary Figure 2**). We believe AIM2 might undergo slight oligomerization in the absence of dsDNA prior to and during the single-molecule experiments, even though AIM2 elutes as a monomer at 10 – 14 nM during the last protein purification step (**Supplementary Figure 3**, Methods). It has been reported using ns-TEM (Transmission Electron Microscopy) that AIM2 polymerizes into filaments in the absence of DNA at a concentration greater than 500 nM ^27^. After extensive ns-TEM analysis of our AIM2 solutions at ∼ 10 nM, we were not able to clearly observe filaments likely due to the low concentration. The use of single-molecule techniques might have facilitated the detection of these assemblies.

### AIM2-DNA dissociation rate constant (k_off_) at the single-molecule level

Experiments using correlative confocal fluorescence microscopy and optical tweezers allow to analyze the association and dissociation kinetics between AIM2 and the single dsDNA molecule (**Figure 4a**). Specifically, we determined the time single AIM2 molecules and AIM2 self-assemblies remain bound to the dsDNA molecule. For this purpose, the fluorescence intensity (photon counts) of fluorophore tagged AIM2 is recorded as a function of time and genomic position in kymographs. **Figure 4b** shows a representative sample of typical kymographs obtained for AIM2 oligomers of different sizes with traces of different intensity and retention times. To separate AIM2 oligomerization from DNA binding, we selected hundreds (N = 316) of single-molecule traces to obtain residence times (**Figure 4c, d**). These traces correspond to an average of 10 photon counts, thus in good agreement with the 11 photon counts determined by the photodepletion step analysis (Methods, **Supplementary Figure 1b**). The residence time distribution of the single-molecule traces was used to determine the AIM2-DNA dissociation rate constant (k_off_) by fitting the histogram to an exponential equation (**Figure 4d**). The k_off_ for single AIM2 molecules to dissociate from the dsDNA molecule is 0.29 ± 0.01 s^-1^. To our knowledge, this is the first dissociation rate constant obtained for a dsDNA inflammasome sensor. This value is approximately two orders of magnitude larger than typical values reported for transcription factors^39,40^. A higher tendency of inflammasome DNA-sensors to detach from DNA could be expected based on the known lack of sequence specificity^25^. Interestingly, similar k_off_ values have been observed for the Klenow fragment of *E. coli* polymerase I (0.40 ± 0.01 s^-1^), which does not require sequence specificity for binding to the template DNA^41^.

**Figure 4.**
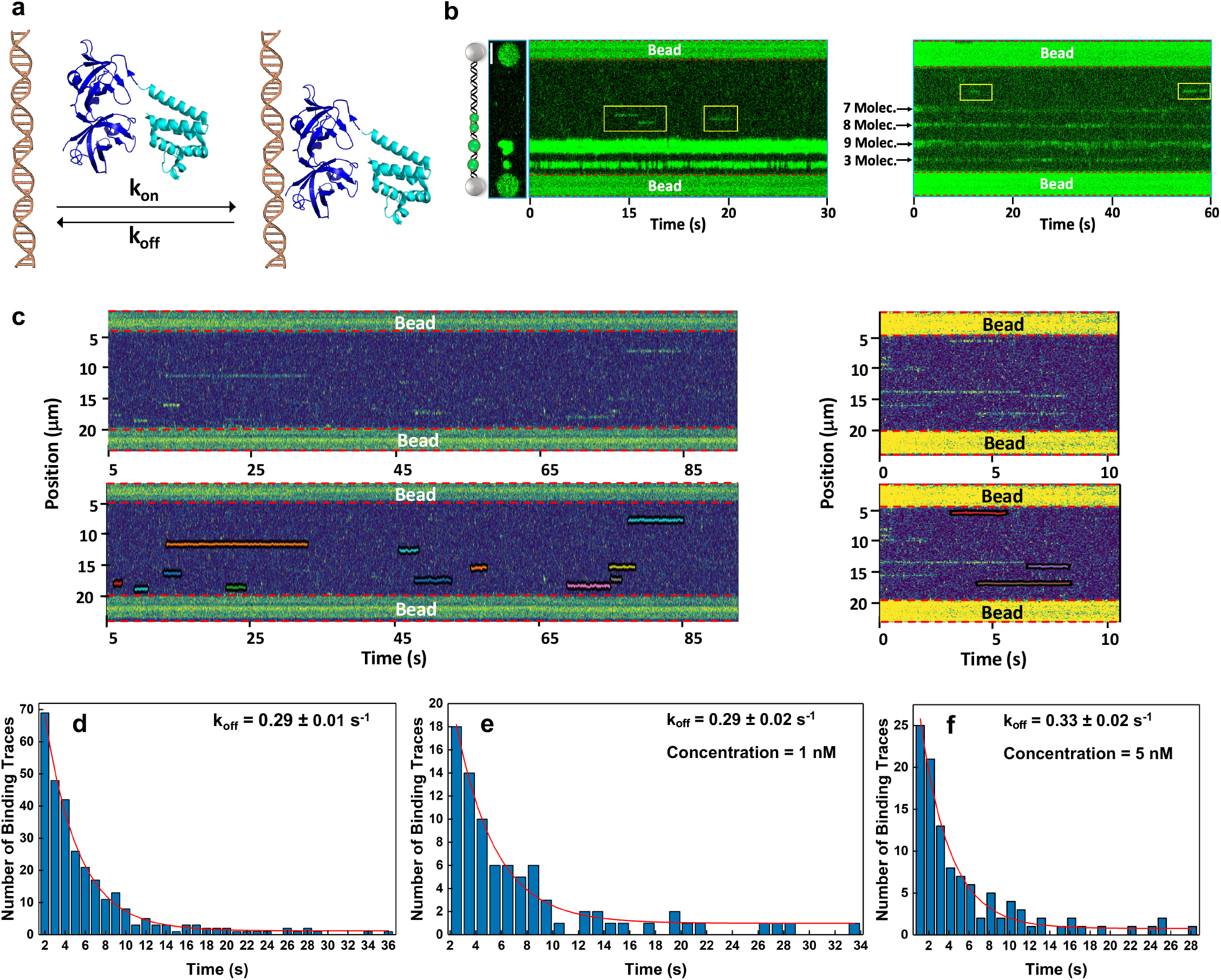
Kinetics of the association and dissociation of single AIM2 molecules to dsDNA. **a** Schematic representation of the association and dissociation of AIM2 to dsDNA (shown at different scales) and the corresponding rate constants. The structures of the PYD^48,49^ and HIN^25^ domains of AIM2 are shown in cyan and dark blue, respectively. **b** Examples of two kymographs showing the coexistence of traces corresponding to large clusters and single-molecule traces (kymograph at the left), and small clusters composed of different numbers of protomers (kymograph at the right). Single-molecule traces are encompassed by yellow boxes. **c** Examples of kymographs (top) and the resulting single-molecule trace tracking (bottom). Only traces appearing after the kymograph started are selected to avoid biases in permanence time determination (kymograph at the right). **d, e, f** Dwell time analysis of AIM2 permanence on dsDNA at different concentrations (d), 1 nM (e) and 5 nM (f) concentration. The red lines represent the fittings to a single exponential function reporting the dissociation rate constant (k_off_). The goodness of fit is represented by R-square and RMSE (root mean squared error) values of 0.99, 0.96, 0.97 and 1.5, 1.0, 1.1 for the three fittings at all concentrations, 1 nM and 5 nM, respectively.

To determine the potential effect of protein concentration on the dissociation rate, we have selected approximately 100 single-molecule traces in kymographs acquired at 1 nM and 5 nM (**Figure 4e, f**). The obtained k_off_ values at the two concentrations are 0.29 ± 0.02 s^-1^ and 0.33 ± 0.02 s^-1^, respectively. These values are similar to the k_off_ obtained with the 316 traces at different concentrations (0.29 ± 0.01 s^-1^), thus indicating that the dissociation rate of the AIM2-DNA complex does not depend on the protein concentration as expected.

We have observed that AIM2 clusters composed of 3 or more protomers are permanent and thus remain attached to the dsDNA molecule for the total length of the kymographs, longer than 20 minutes in some instances (**Supplementary Table 1, Figure 4b**). These results indicate that self-assembly is critical to modulate the k_off_ of AIM2-dsDNA complexes. Altogether, the dissociation data of AIM2-DNA support the concept of inflammasome formation being almost an irreversible process^29^ for sufficiently large oligomers.

### AIM2-DNA association rate constant (k_on_) at the single-molecule level

Real-time fluorescence anisotropy measurements in bulk have been used to determine binding rates of full-length AIM2 and ds-DNA^29^. These thorough studies report that full-length AIM2 assembles into 600-bp dsDNA with an observed rate of ∼ 0.8 min^-1^ (at 72 nM AIM2)^29^. The observed rate in bulk includes both k_on_ and k_off_; however, we can assume that the k_off_ is zero for the longest DNA (permanent oligomer), thus resulting in a k_on_ of 0.18·10^6^ M^-1^ s^-1^ at the reported AIM2 concentration (**Table 1**).

**Table 1:** Association and dissociation rates and equilibrium dissociation constant (K_D_) of AIM2 and IFI16 variants to dsDNA

| | $k_{\text{off}}$ ($\text{s}^{-1}$) | $k_{\text{on}}$ ($\text{M}^{-1} \text{s}^{-1}$) | $K_D$ (nM) |
| --- | --- | --- | --- |
| Single molecule: AIM2<sup>FL</sup> | $0.29 \pm 0.01$ | $0.37 \cdot 10^6 - 1.81 \cdot 10^6$ | 160 - 780 |
| Fluorescence Anisotropy: AIM2<sup>FL</sup> | N/A | $0.18 \cdot 10^6$<sup>a</sup> | $\leq 3$ |
| Fluorescence Anisotropy: AIM2<sup>HIN</sup> | N/A | N/A | $176 \pm 35$<sup>b</sup> |
| Fluorescence Anisotropy: AIM2<sup>HIN</sup> | N/A | N/A | $212 \pm 28$<sup>c</sup> |
| Fluorescence Anisotropy: MBP-AIM2<sup>FL</sup> | N/A | N/A | $234 \pm 42$<sup>c</sup> |
| Fluorescence Anisotropy: MBP-AIM2<sup>HIN</sup> | N/A | N/A | $584 \pm 22$<sup>c</sup> |
| Fluorescence Anisotropy: IFI16<sup>FL</sup> | N/A | N/A | $65 \pm 19$<sup>d</sup> |

<sup>a</sup> $k_{\text{on}}$ extracted from reported $k_{\text{obs}} \sim 0.8 \text{ min}^{-1}$ using FAM-dsDNA600, 72 nM AIM2<sup>FL</sup> and 160 mM KCl<sup>29</sup>
<sup>b</sup> FAM-dsDNA20 (200 mM NaCl)<sup>25</sup>
<sup>c</sup> FAM-dsDNA72 (160 mM KCl)<sup>27</sup>
<sup>d</sup> FAM-dsDNA72 (160 mM KCl)<sup>26</sup>

To determine the association rate (k_on_) of full-length AIM2 on the single λ-dsDNA, we analyzed over 100 single-molecule traces in kymographs acquired with a constant time length of 600 s. Importantly, the observed traces do not show attachment and detachment in the same genomic position, which is expected based on the absence of sequence specificity. Therefore, it has not been possible to identify “unbound” periods (t_on_) between traces of protein attached to the same position in the DNA, thus making this measurement challenging. Therefore, we have considered the “unbound” period as the total observation time minus the sum of the residence time of all traces for each individual kymograph (**Supplementary Tables 2, 3**). This analysis has been done at 1 nM and 5 nM starting protein concentration values.

We noticed kymographs acquired at 1 nM with more than 20 single-molecule traces, whereas most kymographs acquired at 5 nM show 11-14 traces (**Supplementary Tables 2, 3**). This observation suggests that increasing the AIM2 concentration decreases the number of single molecules due to oligomerization in the absence of dsDNA, thus influencing the t_on_ values. Therefore, it is more reliable to report a range of association rate constant values (k_on_) obtained at the two concentrations (**Table 1**). This interval agrees overall with the k_on_ derived from the assembly rate determined by the fluorescence anisotropy experiments in bulk^29^ (**Table 1**).

### Dissociation constant (K_D_) determination of single AIM2-DNA complex devoid of effects from AIM2 oligomerization

Using fluorescence anisotropy in bulk, the dissociation constant (K_D_) between full-length AIM2 and DNA has been estimated to be ≤3 nM^27^. However, the K_D_ increases to 212 ± 28 nM for the truncated construct lacking the PYD (AIM^HIN^)^27^. A similar K_D_ value (K_D_ = 176 ± 35 nM) has been reported for AIM2^HIN 25^ (**Table 1**). The K_D_ values in the presence and absence of the PYD suggest that higher affinity requires protein oligomerization via PYD. However, truncation of the protein could result in unwanted structural and functional modifications. Thus, separating protein-DNA binding from protein oligomerization using the native sequence will report on the affinity of AIM2 for DNA devoid of effects from protein oligomerization.

The estimated k_on_ values and the determined k_off_ using single-molecule analysis allow an approximation of the range of K_D_ values for the AIM2-DNA complex (**Table 1**). These K_D_ values are more than three orders of magnitude larger than the K_D_ reported based on the fluorescence anisotropy studies for full-length AIM2^27^, and close to the values reported for the truncated AIM2 lacking the PYD domain^25,27^. This comparison highlights that the K_D_ values of full-length AIM2 at the single-molecule level are not affected by protein oligomerization (**Table 1**). Therefore, the single-molecule results separate the effect of protein-self association from DNA binding and indicate that the affinity of native AIM2 for dsDNA falls in the sub-micromolar range in the absence of oligomerization.

### DNA-bound AIM2 oligomers increase size fourfold in seconds

Using real-time FRET (Förster Resonance Energy Transfer) and fluorescence anisotropy in bulk, it has been reported that assembly rates of full-length AIM2 and DNA depend on the length of the latter^27,29^. An increase in assembly rates close to 700-fold has been observed when extending dsDNA from 24-to 600-bp^29^. However, we lack information on oligomer size and oligomer growth rates in the presence of sufficiently long DNA.

To obtain this information, we have monitored the growth rate of AIM2 clusters bound to λ-phage dsDNA (48.5 kbp). The change in photon counts as a function of time reports on the growth of oligomers with different starting numbers of molecules: 1, 2 and 4 molecules (**Figure 5a, b, c**). In some instances, the oligomers can quadruplicate their size in approximately 4 s (**Figure 5a, b**). The lack of steady increase in photon counts is likely due to fluorophore blinking; however, a clear upward trend is observed. Oligomer growth rates, obtained as the difference between final and initial photon counts divided by the total observation time, tend to increase with AIM2 concentration as expected: 7.4 s^-1^, 10.7 s^-1^ and 34 s^-1^ at 0.5 nM, 2 nM and 13.5 nM, respectively.

**Figure 5.**
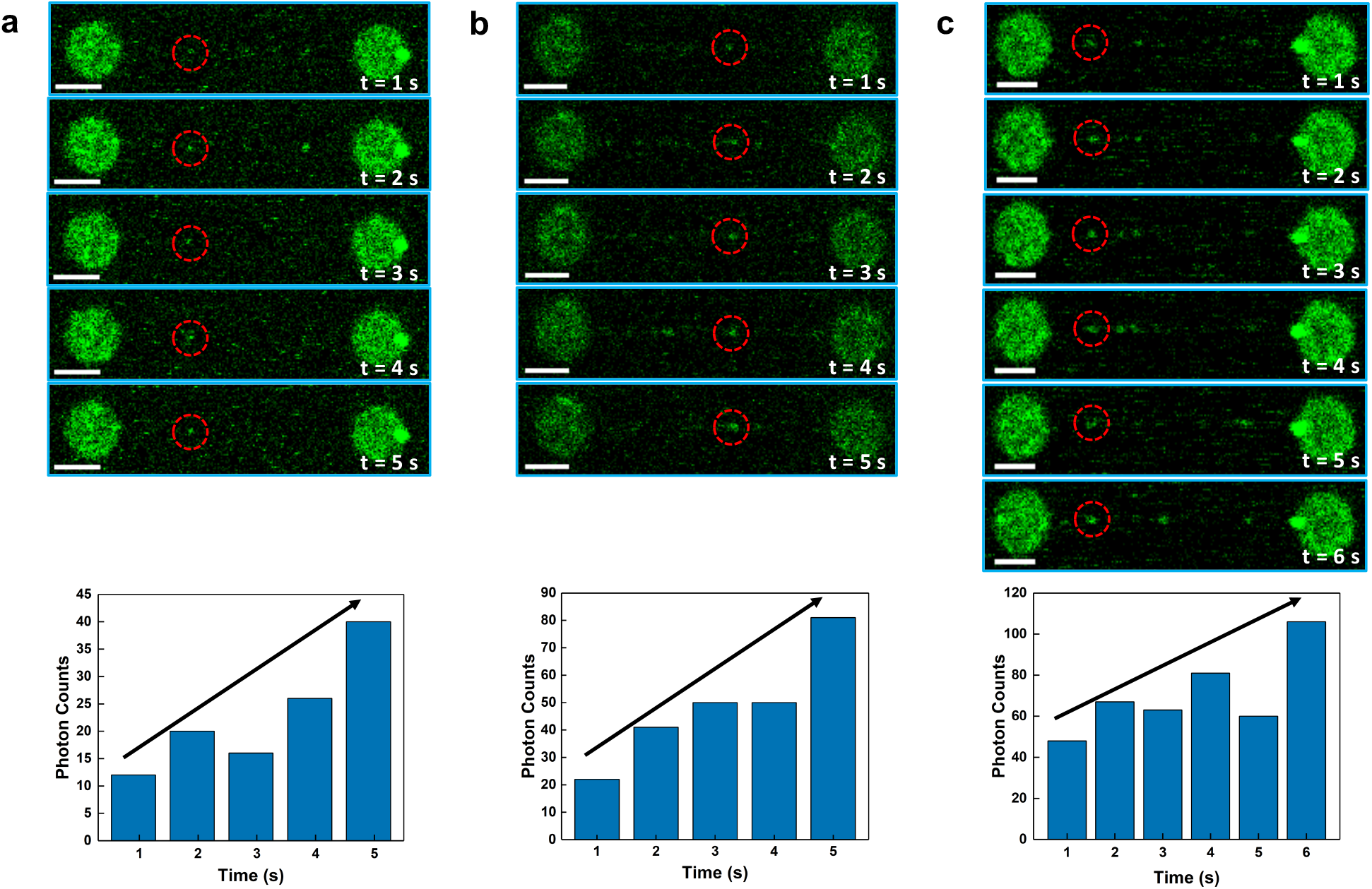
Growth rate of AIM2 oligomers bound to dsDNA. Top: Two-dimensional frames of confocal scans indicating AIM2 binding to the single dsDNA acquired at the times indicated in white. Starting number of molecules in the fluorescent spot circled in red are **a** 1; **b** 2 and **c** 4. Scale bar in white represents 3 µm. Bottom: Overall increase in photon counts with time for the three fluorescent spots corresponding to 1, 2 and 4 starting molecules.

### DNA-bound oligomers grow via distinct mechanisms

Two-dimensional frames of confocal images captured in continuous scanning mode were analyzed to identify specific cluster growth directions. Average pixel brightness (gray values) calculated along the vertical axis of the 2D frame are represented as a function of distance along the DNA molecule resulting in intensity profile plots. The comparison of profile plots from 2D frames acquired at different times shows that AIM2 oligomers grow along both left and right directions of the dsDNA (**Figure 6a, b**). This observation implies that oligomer growth happens by incoming AIM2 molecules binding to the dsDNA at either side of a particular cluster or previously bound molecule. This is an expected result considering that AIM2 does not show DNA sequence specificity and suggests that AIM2 binding to either side of a pre-existing oligomer is stochastic. Interestingly, we have observed an increase in fluorescence intensity not localized at either side of the dsDNA (**Figure 6c)**, indicating that oligomer growth can happen by incoming AIM2 molecules not binding directly to the dsDNA but instead by binding to AIM2 molecules already forming part of the oligomer (**Figure 6c**). Since the function of the PYD in AIM2 is to participate in protein-protein interactions, this type of oligomer growth is likely driven by PYD-PYD binding and does not depend on interactions with the DNA. This type of oligomer would expose AIM2 molecules with free HIN domains available to interact with other dsDNA molecules or fragments. The possibility of AIM2 oligomers growing by incorporating the incoming AIM2 molecules independently of DNA binding explains the observation of asymmetric clusters (**Figures 2b, 4b**).

**Figure 6.**
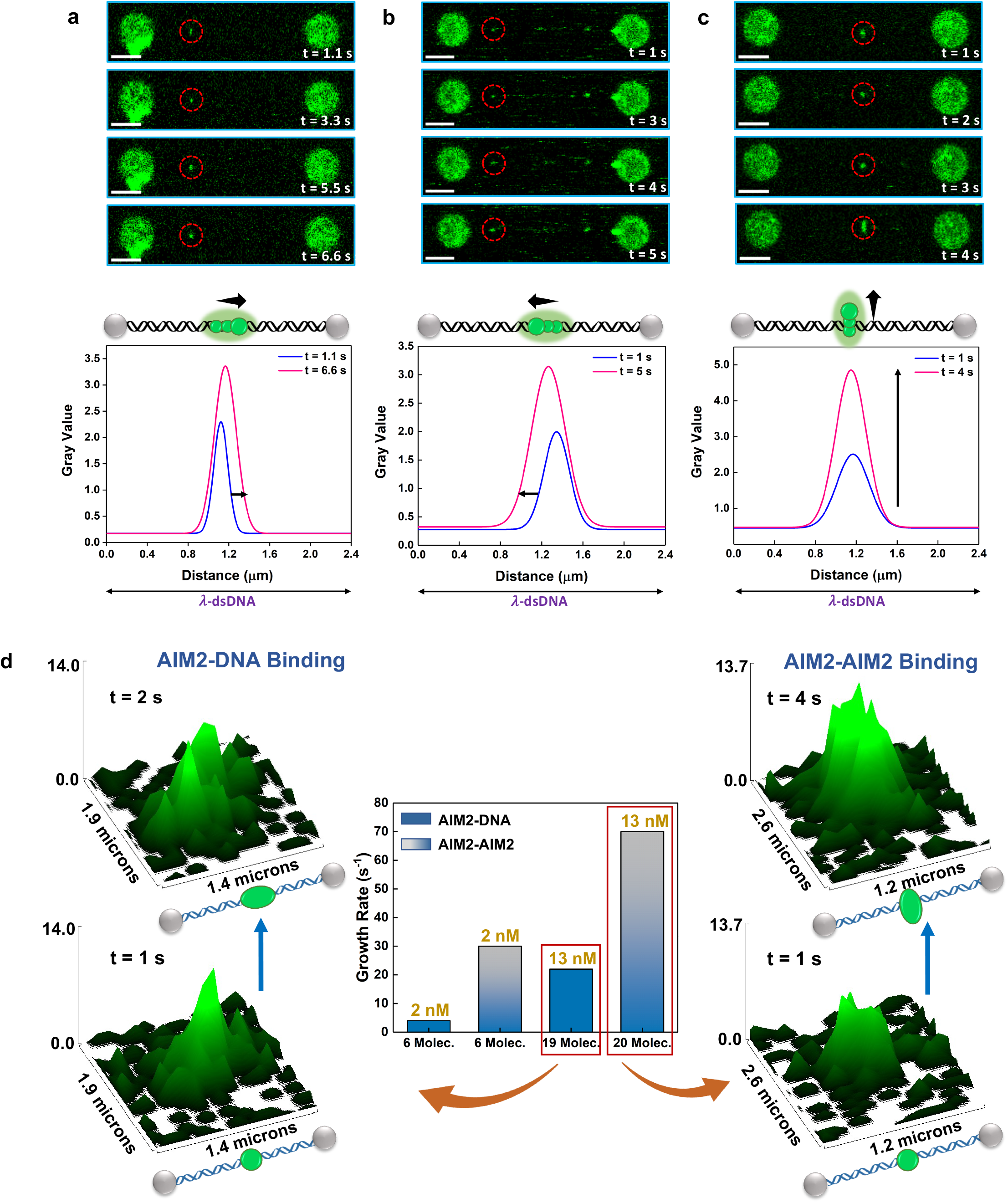
AIM2 oligomers grow via two distinct mechanisms. **a, b, c** Top: Two-dimensional scans of three AIM2 oligomers (circled in red) growing as a function of time. Bottom: Oligomer growth represented by the increase in fluorescence intensity (gray value) to the right (a) and left (b) of the DNA molecule and in direction perpendicular to the DNA (c). **d** Center: Oligomer growth rates as determined by changes in fluorescence intensity with time for clusters with different starting number of molecules (numbers in the X-axis in black) at two different AIM2 concentrations (numbers in gold on top of the bars). The different bar colors represent the two mechanisms of oligomer growth via AIM2 binding to DNA (blue) and AIM2-AIM2 binding in the absence of DNA binding (blue-gray). Higher growth rates are observed for the AIM2-AIM2 binding mechanism at both concentrations. Left and right: Three-dimensional plots derived from AIM2 oligomers with 19 and 20 molecules, representing changes in fluorescence intensity as a function of time (time increasing from bottom to top) along the DNA axis (indicated by schematic representation of DNA and trapped beads) and in direction perpendicular to the DNA. Growth via AIM2-DNA binding (left) shows increased fluorescence along the DNA axis. Growth via AIM2-AIM2 binding (right) shows increased fluorescence along the axis perpendicular to the DNA.

Furthermore, the growth rate is faster for oligomers increasing through PYD-PYD binding compared to clusters with incorporation via DNA binding at the same protein concentration and similar starting number of molecules. **Figure 6d** shows significantly different growth rates for two clusters with 19 and 20 molecules, respectively. The slower rate corresponds to a cluster with AIM2 incorporation along the DNA, as shown by the increase in fluorescence intensity in this direction. In contrast, the oligomer that grows faster has the incorporation of AIM2 molecules in direction perpendicular to the DNA, as shown by the increase in fluorescence in this direction (**Figure 6d**).

### AIM2 does not diffuse along the dsDNA molecule

The interaction between IFI16 and dsDNA has been studied previously by TIRF^28^. IFI16 molecules diffuse several µm along λ-phage dsDNA to find already bound IFI16 clusters for interaction^28^ (**Figure 7a**). Diffusion decreases and stops when clusters grow to ∼ 8 molecules^28^. It was found that the minimum dsDNA length required for efficient IFI16 oligomerization is 50-70 bp. The presence of nucleosomes results in shorter dsDNA sequences, thus hindering IFI16 diffusion and self-assembly ^28^ (**Figure 7a**). In contrast, low chromatinization of foreign dsDNA leads to the exposure of long stretches of dsDNA, thus allowing IFI16 to diffuse freely and oligomerize. Based on these results, the authors elegantly explain how IFI16 discriminates between host and foreign dsDNA^28^.

**Figure 7.**
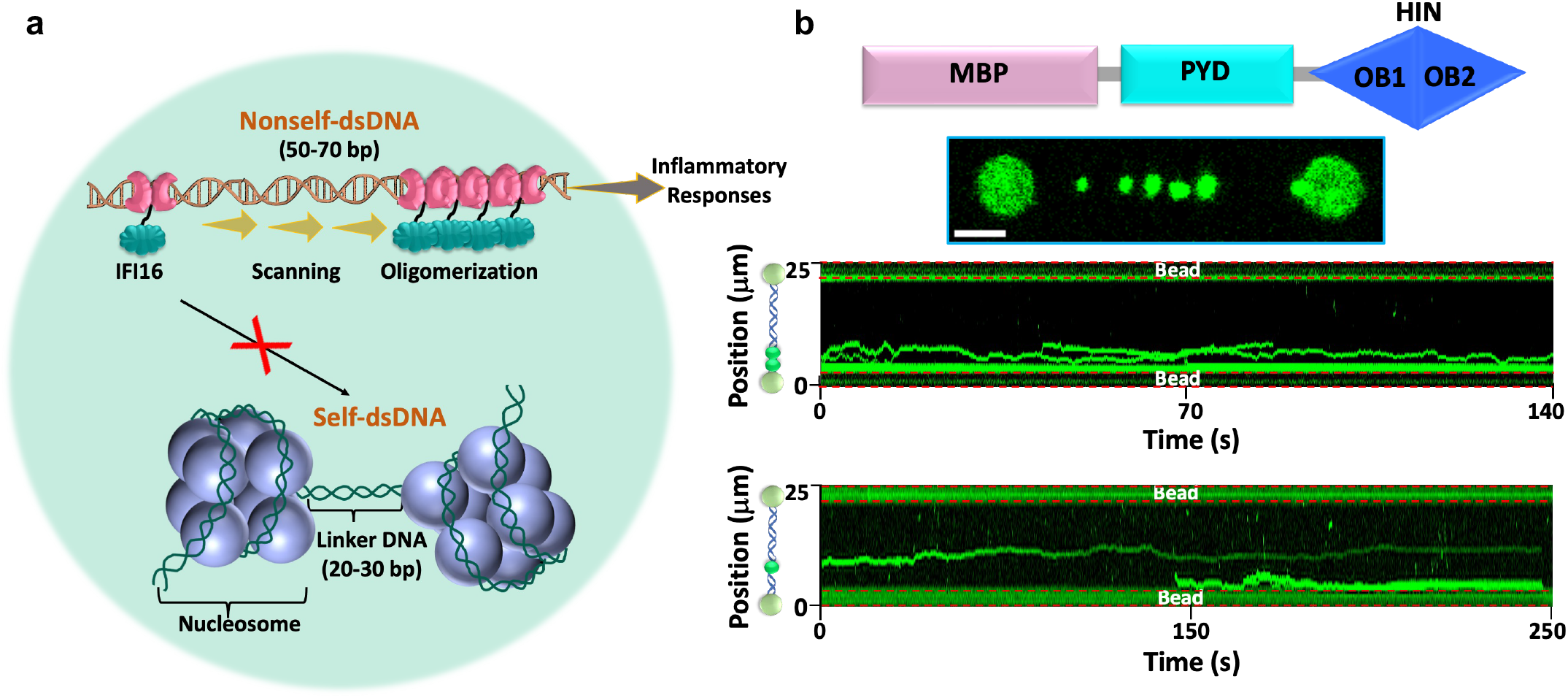
MBP-AIM2 diffuses along the dsDNA analogously to IFI16. **a** Schematic representation of IFI16 diffusion along the dsDNA as a mechanism to discriminate between self- and foreign-DNA^28^. **b** Typical two-dimensional scan (680 pM MBP-AIM2 and 160 mM KCl) and kymographs (1 nM and 680 pM MBP-AIM2, top and bottom, respectively, and 160 mM KCl) showing dsDNA binding and diffusion of MPB-AIM2.

We show here that AIM2 behaves differently as it does not diffuse along the dsDNA (**Figure 4b, c**). Once attached to a certain genomic position, both AIM2 clusters and single molecules do not change position during long and short permanence times, respectively. At the protein domain level, AIM2 contains only one dsDNA binding domain, whereas IFI16 bears two (**Figure 1b, d**). The presence of two HIN domains in IFI16 could explain the different permanence times and diffusivity between the two proteins. In fact, the TIRF study reports that a truncated IFI16 construct lacking the PYD shows analogous diffusion behavior^28^, which points to the key role of the HIN domains in the DNA binding mode.

The different behavior shown by AIM2 is intrinsic to the protein and not related to the experimental conditions. In fact, we have been able to observe diffusion along the dsDNA for a construct of AIM2 carrying the MBP (Maltose Binding Protein) tag (**Figure 7b and Supplementary Figure 4**). The capability of MBP-AIM2 to diffuse along the DNA must be related to the presence of the tag. It has been reported previously that the binding affinity of MBP-AIM2 for dsDNA is at least two orders of magnitude smaller than that of untagged AIM2 (**Table 1**), likely due to MBP interfering with the PYD-PYD driven oligomerization^27^. However, our results on the absence of diffusion of single AIM2 molecules do not include a PYD-PYD binding effect, thus indicating that the MBP tag could directly affect the interaction between the HIN domain and DNA, which could lead to diffusion. Binding affinity values of AIM2 constructs lacking the PYD are a better reference for our single-molecule data because the latter are not affected by oligomerization. For instance, the reported K_D_ of MBP-AIM2^HIN^ for dsDNA is 2.7 times larger than that of AIM2^HIN^ (lacking the PYD and the MBP tag) (**Table 1**)^27^, which points to the effect of the MBP tag on the binding between the HIN domain and dsDNA. Overall, our results on the diffusion of MBP-AIM2 and the comparison with the previously reported K_D_ values indicate that a stronger interaction between DNA and AIM2 likely explains the absence of diffusion.

## Discussion

Detailed studies on the operating mode of the inflammasome sensor AIM2 have revealed that dsDNA binding and protein oligomerization are connected and cooperative^27^. Compelling evidence support a mechanism in which the dsDNA acts as a platform for AIM2 oligomerization^25,27^. In addition, it has been shown that filaments formed by AIM2 oligomerization upon dsDNA binding function as a template for the polymerization of the inflammasome adaptor ASC^27^. This mechanism helps explain the robust inflammatory response observed upon cell treatment with dsDNA fragments sufficiently long to trigger the formation of the AIM2 inflammasome^25^.

We have presented here fundamental information on the kinetics and mechanism of propagation of AIM2-DNA assemblies to further our understanding of the initial stages of inflammasome formation at the molecular level. AIM2 oligomerizes into distinct, individual assemblies of different sizes coexisting on the same dsDNA molecule (**Figure 2b**). Small oligomers composed of 1-20 molecules grow with a rate ranging from 0.2 - 6 molecules per second at low nanomolar concentrations. This time scale aligns with previously reported kinetic studies in live cells showing that the ASC speck assembles in approximately 3 minutes once ASC concentration redistributes upon AIM2 inflammasome activation^42^. The long-lived nature of the ASC speck is demonstrated in these studies by its persistence for several hours^42^. Our results on the permanent attachment of AIM2 oligomers are in accord with these observations.

Based on the association and dissociation rates of AIM2 and dsDNA at the single-molecule level, we have determined the affinity of AIM2 for dsDNA devoid of self-association effects. The K_D_ falls in the sub-micromolar range, likely due to the lack of sequence specificity for dsDNA. As expected, AIM2-DNA affinity is significantly low (approximately two orders of magnitude) compared to sequence-specific proteins such as transcription factors.

We have shown here that AIM2 oligomerizes to some extent in the absence of dsDNA at sub-nanomolar concentration. The use of single-molecule techniques likely has facilitated the detection of these assemblies. AIM2 self-association unaccompanied by dsDNA binding could lead to sterile inflammation. A potential mechanism to tightly control AIM2 self-association could involve keeping a low basal concentration. In this regard, our results suggest that AIM2 basal concentration is likely lower than sub-nanomolar. Additionally, regulation of self-association could depend on inhibitory mechanisms. In fact, reported evidence on the interaction between the PYD and HIN domains of AIM2 suggest an autoinhibitory mechanism when AIM2 is in a resting state^25^. It is possible that autoinhibition competes with oligomerization and dsDNA binding at very low basal concentrations. Based on our results and in accordance with previous reports, we hypothesize that once pathogenic dsDNA enters the cell cytoplasm, AIM2 concentration increases, thus facilitating protein oligomerization and irreversible dsDNA binding, which subsequently leads to active inflammasome formation and downstream signaling until the cell dies.

Finally, we have shown here that AIM2 does not diffuse along the dsDNA. Unlike the nuclear sensor IFI16, AIM2 does not need to distinguish between host and foreign dsDNA, as only aberrant host dsDNA can be found in the cytosol. Thus, AIM2 does not need to diffuse on sufficiently long stretches of dsDNA to facilitate oligomerization, which is the proposed mechanism for IFI16 to discriminate between host and foreign dsDNA^28^. We hypothesize that specific interactions of the HIN domains of AIM2 and IFI16 with dsDNA, as well as the presence of one versus two HIN domains, lead to the different dsDNA binding behavior. In fact, the reported binding affinity between full-length IFI16 and dsDNA is significantly smaller than that of AIM2 (**Table 1**)^26,27^. Another discrepancy in the behavior of AIM2 and IFI16 is that the HIN domain of AIM2 has been shown to oligomerize in the DNA^27^, unlike the HIN domains of IFI16^26^. Moreover, the X-ray structures of the complexes between AIM2-HIN and IFI16-HINb with DNA revealed that the former leads to a larger solvent-accessible surface area buried upon complex formation^25^.

Based on the data reported here, we suggest a model for the formation of AIM2-DNA assemblies that involves three complementary and likely simultaneous scenarios (**Figure 8**). Single AIM2 molecules stochastically and transiently bind to dsDNA with low affinity leading to survival times of approximately 3 s (**Figure 8a**). During this time, incoming AIM2 molecules bind to dsDNA in positions that are sufficiently close to prebound AIM2 molecules or oligomers, causing lateral growth of the oligomer and permanent attachment (**Figure 8b**). In conjunction or alternatively, incoming AIM2 molecules may also bind to prebound clusters via AIM2-AIM2 interactions (in the absence of dsDNA binding), leading to oligomer growth in directions perpendicular to the dsDNA (**Figure 8c**). Overall, AIM2-DNA assembly formation and propagation can be understood as a polymerization process with primary (AIM2-DNA binding) and secondary (AIM2-AIM2) nucleation events. The double-nucleation mechanism is a well-known process demonstrated previously for sickle fiber formation by hemoglobin S polymerization^43^. The different propagation mechanisms proposed here (**Figure 8b, c**) likely favor the formation of intertwined and densely packed filamentous structures via 1) AIM2^PYD^-AIM2^PYD^ interactions between nucleoprotein filaments; 2) AIM2^HIN^ interactions between nucleoprotein filaments and free dsDNA fragments; and 3) AIM2^PYD^-ASC^PYD^ interactions between nucleoprotein filaments and ASC or polymerized ASC. This mechanism explains the random appearance of AIM2 clusters of different sizes and shapes (**Figure 2b, 4b**), likely resulting from the favored binding of incoming AIM2 molecules to larger pre-existing bound oligomers. Additionally, attached AIM2 clusters of increasing size limit dsDNA availability, thus leading to a higher probability of incoming molecules to interact with prebound oligomers.

**Figure 8.**
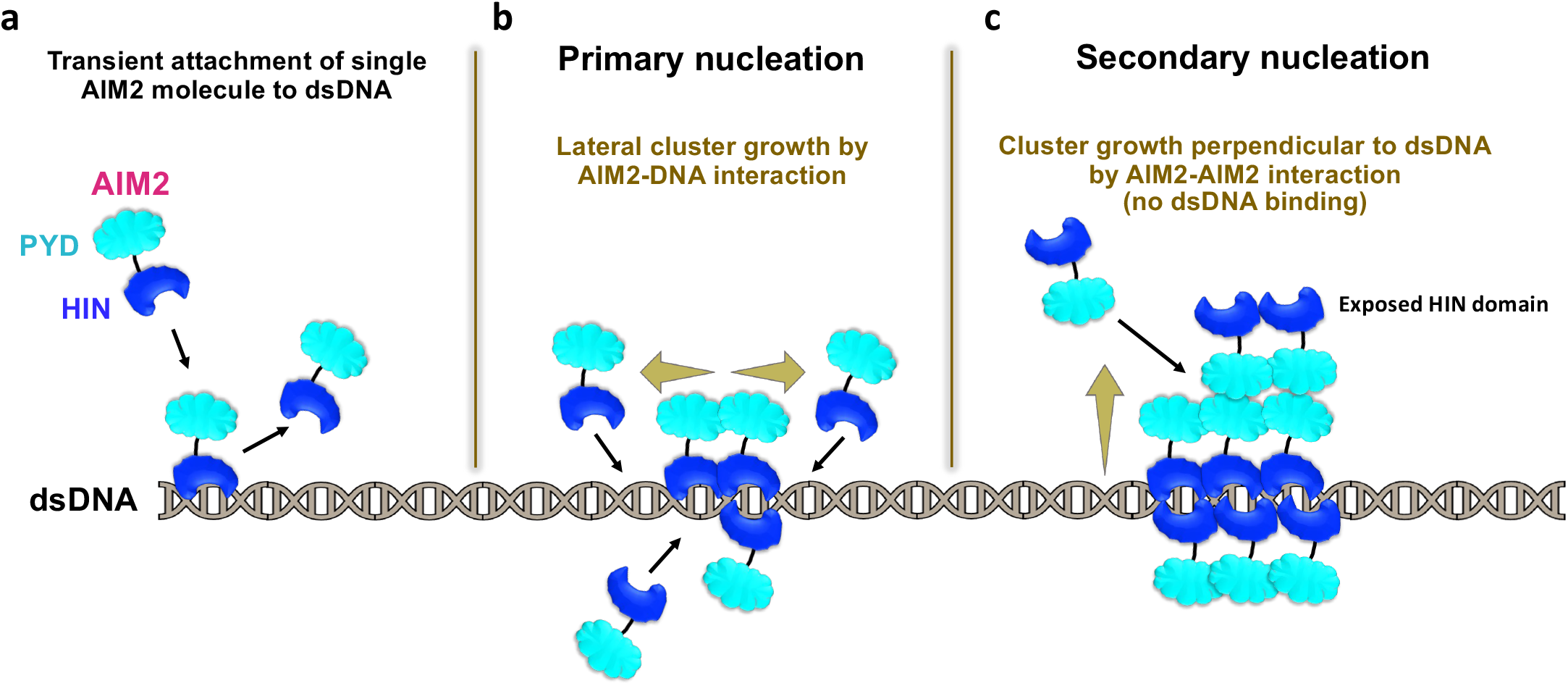
Primary and secondary nucleation steps in the assembly mechanism of the AIM2 inflammasome sensor. **a** Stochastic attachment of AIM2 to dsDNA and detachment. **b** Attachment of AIM2 molecules to prebound AIM2 oligomer via dsDNA binding, resulting in permanent attachment. Cluster grows along the dsDNA. **c** Attachment of AIM2 molecules to prebound AIM2 oligomer via AIM2-AIM2 interaction in the absence of dsDNA binding, resulting in permanent attachment. Cluster grows in direction perpendicular to dsDNA.

## Methods

### Synthesis and cloning of human AIM2

The full-length human AIM2 gene (amino acids 1-342) with an N-terminal 6xHis tag followed by a MBP tag and Tobacco Etch Virus protease (TEVp) recognition site (ENLYFQG) was synthesized and cloned into the pET21(b) vector by Gene Universal Inc. In addition, this construct includes a sortase A recognition site (LPETG) connected to the C-terminus of AIM2 by a flexible linker (GGGGS) and with two extra glycine residues after the recognition site to ensure optimal results of the sortase-mediated transpeptidation reaction^44^ that is used to label AIM2 with a fluorophore-tagged peptide.

### Expression and purification of AIM2 constructs

The AIM2 construct was transformed into Rosetta BL21 (DE3) cells, which were grown overnight in LB medium supplemented with 100 µg/mL ampicillin and 34 µg/mL chloramphenicol at 37 °C and 220 rpm. Overnight seed culture was transferred into large volume of LB medium and grown at 37 °C to an OD_600_ of 0.6-0.8. Protein expression was induced at 18 °C with 1 mM isopropyl β-D-1 thiogalactopyranoside (IPTG) and incubated overnight. Cells were harvested and resuspended in lysis buffer containing 20 mM HEPES, pH 7.4, 400 mM KCl, 1 mM BME, 0.1% Triton X-100 and 5% glycerol. The cells suspension, supplemented with 100 mM phenylmethylsulfonyl fluoride (PMSF), 100 µg/mL lysozyme, and a protease inhibitor cocktail (Pierce protease inhibitor tablet contains AEBSF, aprotinin, bestatin, E-64, leupeptin, and pepstatin), was incubated at 4 °C for 30 min. Cells were lysed by at least 7 cycles of freeze-thaw using dry ice/ethanol bath and centrifuged at 35,000 rpm for 40 min. The supernatant was collected, and the pellet was washed again in lysis buffer. Supernatants were collected after each centrifugation step. To remove the bacterial DNA, a 2-3% solution of streptomycin sulfate was slowly added into the supernatant, followed by constant stirring at 4 °C for 30 min. The insoluble precipitates were separated by centrifugation at 35,000 rpm for 40 min. The streptomycin sulfate precipitation step was repeated twice to achieve efficient elimination of DNA.

The supernatant obtained after centrifugation was filtered with a 0.45 µm pore filter and applied onto a prepacked 5 mL Ni-NTA column (Thermo Fisher Scientific) preequilibrated with lysis buffer. The column was washed in two steps with lysis buffer containing 25 mM and 50 mM imidazole (50 mL each). MBP-AIM2 was eluted in lysis buffer containing 400 mM imidazole. Fractions were collected and analyzed by SDS–PAGE. Fractions containing MBP-AIM2 were pooled and purified by ion-exchange chromatography using 5 mL HiTrap-SP column (GE Healthcare) equilibrated in 20 mM HEPES, pH 7.4, 160 mM KCl, 1 mM BME, 0.1% Triton X-100, and 5% glycerol on an HPLC system. After washing, the protein was eluted with a linear gradient ranging from 0.16 M to 1 M KCl at a flow rate of 1 mL/min and subsequently analyzed for purity by SDS–PAGE. Protein concentration was determined by UV absorption measured at 280 nm using a molar extinction coefficient (ε) of 75,290 M^-1^ cm^-1^ for MBP-AIM2. The A_260_/A_280_ ratio of the different fractions was measured to identify DNA-free protein, and the fractions with a ratio of ∼ 0.64 were pooled. Dialysis and concentration steps of protein solutions were avoided throughout the purification process to reduce protein oligomerization.

To remove the MBP tag, 450 nM MBP-AIM2 was mixed with 18 µM TEVp in a buffer containing 20 mM HEPES, pH 7.5, 160 mM KCl, 1 mM BME, 0.1% Triton X-100, and 5% glycerol and incubated at 30 °C, 220 rpm for 1 h. The reaction was subjected to centrifugation to separate protein precipitation due to MBP removal, and the supernatant was further used for fluorescent labeling. For experiments requiring the MBP-AIM2 construct, the TEVp cleavage step was omitted.

### Labeling of AIM2 constructs with Alexa Fluor 488

To ensure single-fluorophore labeling of AIM2 and avoid mutations commonly used in fluorophore labeling strategies, a short peptide conjugated with Alexa Fluor 488 was attached covalently to the protein constructs using sortase A transpeptidation.

#### Labeling of the short peptide with Alexa Fluor 488 maleimide dye

A peptide with amino acid sequence, GGGC, was synthesized and purified by Thermo Fisher Scientific. The peptide was dissolved in a labeling buffer containing 20 mM sodium phosphate, pH 7.2, and 100 mM NaCl. Alexa Fluor 488 C_5_ maleimide (Thermo Fisher Scientific) was dissolved in DMSO. The labeling reaction was prepared by mixing a 300 µM peptide solution and Alexa 488 at a 2-fold molar excess relative to the peptide concentration. The reaction mixture was incubated at 25 °C for 4 h at 220 rpm. The reaction was quenched by adding 2-Mercaptoethanol (BME). The labeled peptide was purified by reverse-phase chromatography using ZORBAX 300SB-C18 column (Agilent) equilibrated with 5% acetonitrile, 95% H_2_O, and 0.1 % TFA and eluted in a gradient created with a buffer containing 5% H_2_O, 95% acetonitrile and 0.1 % TFA. The eluted fractions were collected, and labeling efficiency was determined to be ≥ 80% using mass spectrometry. The concentration of the labeled product was determined from the absorbance at 493 nm using a molar coefficient of 72,000 M^-1^ cm^-1^ for Alexa 488. The labeled peptide was lyophilized and stored at -80 °C for further use.

#### Pairing of peptide-Alexa 488 conjugate to MBP-AIM2 and AIM2 (without MBP tag)

The covalent attachment of the GGGC peptide-Alexa 488 conjugate with the LPETG-containing MBP-AIM2 and AIM2 (without MBP tag) was performed by sortase-mediated C-terminal transpeptidation.

To label MBP-AIM2, the protein and the peptide conjugate were mixed at a 1:4 molar ratio. Specifically, solutions of MBP-AIM2 and peptide-Alexa 488 conjugate were mixed in sortase reaction buffer (20 mM HEPES, pH 7.5, 160 mM KCl, 1 mM BME, 0.1% Triton X-100, and 5% glycerol, and 10 mM CaCl_2_), resulting in final protein and peptide concentrations of ∼ 20 µM and 80 µM, respectively. Sortase A was added to a final concentration of 10 µM, and the reaction was incubated overnight in the dark at 4 °C. Subsequently, MBP-AIM2 labeled with peptide-Alexa 488 was purified by size exclusion chromatography (SEC) using Superdex 200 increase 10/300 GL column (GE Healthcare) in a buffer containing 20 mM HEPES, pH 7.5, 160 mM KCl, 1 mM BME, 0.1% Triton X-100 and 5% glycerol buffer at a flow rate of 0.5 mL/min. The concentration of the labeled fraction was determined from the absorbance values at 280 nm and 493 nm by considering the dye’s absorbance at 280 nm using a correction factor (CF) provided by the manufacturer of 0.11. A theoretical molar extinction coefficient of 75,290 M^-1^ cm^-1^ was used for MBP-AIM2 to determine the protein concentration using the equation provided by the manufacturer:

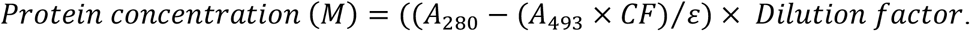

To label AIM2 (without MBP), the protein and the peptide conjugate were mixed in the sortase reaction buffer, resulting in final concentrations of < 450 nM and 5 µM, respectively. Sortase was added to a final concentration of 10 µM. The reaction was incubated overnight in the dark at 4 °C.

The purification protocol was identical to that of MBP-AIM2 (above). Fractions corresponding to monomeric AIM2 were pooled. The use of a short peptide with poly-Gly sequence (GGGC) and a flexible linker (GGGGS) connecting AIM2 to the sortase recognition site match the requirements to achieve ∼ 95% transpeptidation efficiency^44^. The concentration of SEC fractions was determined by fluorescence spectroscopy experiments performed at 25 °C using a Horiba PTI QuantaMaster 400 fluorimeter with a slit width of 10 nm for both excitation and emission. Briefly, fluorescence emission spectra of the labeled protein fractions were collected from 490 nm to 700 nm by exciting Alexa 488 at 480 nm. Maximum emission intensity was obtained at 519 nm. The concentration of the AIM2-peptide-Alexa 488 samples was calculated by extrapolating from a calibration curve based on the fluorescence emission of standard samples prepared from the MBP-AIM2-peptide-Alexa 488 in a concentration range from 0.5 nM to 10 nM.

### Single-molecule experiments using dual-trap optical tweezers with confocal fluorescence microscopy

Single-molecule experiments were performed at room temperature on the C-Trap (LUMICKS) system integrated with optical tweezers, confocal fluorescence microscopy and microfluidics, and recorded using BlueLake software (LUMICKS). Experiments were performed in freshly prepared imaging buffer containing 20 mM HEPES, pH 7.5, 160 mM KCl, 1 mM BME, 0.1% Triton X-100, and 5% glycerol supplemented with an oxygen scavenging system (5 mg/mL D-glucose, 20 µg/mL catalase and 100 µg/mL glucose oxidase) and 1 mM Trolox methyl ether filtered with 0.05 µm syringe filter. All the components (beads, DNA, and protein) were prepared in this buffer. A five-channel laminar flow cell (**Figure 2a**) mounted on an automated XY stage consists of 3 parallel channels (channels 1-3) separated by laminar flow and two orthogonal channels (channels 4 and 5). The flow cell was flushed with buffer ∼ 20 min prior to sample addition. Streptavidin-coated polystyrene beads (Spherotech, 0.5% w/v stock) with a diameter of 3.11 µm were diluted 4/1000 in imaging buffer and placed in channel 1. Channel 2 was filled with biotinylated λ-phage dsDNA (48.5 kbp, LUMICKS, Netherlands) at a concentration of 24 pg/µL. Imaging buffer was flowing in channels 3 and 5, and fluorophore-labeled protein, diluted in imaging buffer to the required working concentration, was placed in channel 4.

Two streptavidin-coated beads in channel 1 were optically trapped with a stiffness of ∼ 0.4 pN/nm using a 1064 nm trapping laser. One molecule of λ-phage dsDNA was tethered between the two beads by moving the traps from channel 1 to channel 2 that are subjected to laminar flow. The traps were moved to channel 3 (buffer), where the presence of a single molecule of dsDNA was verified by comparison of the experimental force-extension curve to the built-in Worm-like chain (WLC) model in BlueLake software. The bead-DNA complex was then moved to protein channel 4 and incubated for protein-DNA binding. Flow from channel 4 was kept off during data acquisition. For confocal imaging, fluorophore-labeled AIM2 was excited at 488 nm, and emission was detected using a blue filter from ∼ 500 to 525 nm. Confocal two-dimensional (2D) images were recorded in single and continuous scanning modes. Kymographs were generated via a confocal line scan through the center of the two beads along the dsDNA molecule in continuous mode. Other imaging conditions included 10% laser power, 50 µs/pixel time, and 100 nm pixel size.

### Processing of confocal images and kymographs

Confocal 2D images and kymographs were exported from BlueLake in HDF5 file formats and processed using custom-written scripts in the Pylake python package provided by LUMICKS. 2D images converted into TIFF format were further analyzed using Fiji for cluster size and growth analysis. In Fiji, the RGB images were split to retain data from the blue channel, converted into green for clear visualization, and the color contrast level was adjusted. A custom-written kymotracking script, available at the script sharing platform Harbor (LUMICKS), was used to track the binding traces in the kymographs and extract information on time, photon counts, and position of individual protein molecule/oligomer on the DNA^45–47^.

### Single-step photobleaching analysis

Kymograph binding traces representing AIM2 oligomers bound to dsDNA were tracked as described above and the obtained information was represented as fluorescence intensity (photon counts) decays versus time. These decay plots were carefully analyzed to identify single and multiple photodepletion steps by calculating the average intensity of flat regions before and after observed intensity drops (**Supplementary Figure 1a**). The distribution of single-step photodepletion steps (N = 50) was binned with a width of 3 photon counts to construct a histogram that was fit to a Gaussian function using Origin (OriginLab 2017). Fitting reveals an average of 11 ± 4 photon counts for a single fluorophore (**Supplementary Figure 1b**). This result was used to calculate the number of molecules forming AIM2 clusters.

### Cluster size distribution and cluster growth analysis

Two-dimensional (2D) confocal images were analyzed in Fiji to determine the number of molecules, surface area, and real-time growth of AIM2 clusters (oligomers) bound to λ-phage dsDNA. Individual clusters were encompassed using Fiji’s rectangle tool, and the obtained fluorescence intensity and surface area were plotted with Origin (**Figure 2c and Figure 3**). All the data were corrected by subtracting the average background intensity for the same area of the selection rectangle. The resulting intensity values were divided by 11 (i.e., the number of photons emitted by a single fluorophore) and further corrected for labeling efficiency to obtain the cluster size (**Figure 3a, b**).

For real-time cluster growth analysis, 2D confocal images were captured in continuous scanning mode and converted into individual, time-stamped, sequential frames using LUMICKS Pylake python package (**Figure 5**). The increase in intensity of individual frames was analyzed in Fiji and plotted as a function of time in Origin (**Figure 5**). Cluster growth direction was determined using Fiji by obtaining profile plots of average pixel brightness (gray values) as a function of distance along the DNA axis. The direction of intensity increase matches the direction of cluster growth (**Figure 6a, b, c**). To generate movies, individual frames were combined at a rate of two frames per second (**Supplementary Figure 4**).

Alternatively, oligomer growth rates shown in **Figure 6d** (center) were determined as the difference between photon counts of final and initial frames divided by the total observation time.

For this analysis, 2D frames of 4 clusters were used, including 2 clusters starting with 6 molecules at 2 nM AIM2, and 2 clusters starting with 19 and 20 molecules at 13.5 nM AIM2 (**Figure 6d, center**). The three-dimensional plots shown in **Figure 6d** (left and right) were generated with Fiji from the analysis of two clusters comprising 19 and 20 molecules.

### Dwell time and kinetic analysis

For kinetic analysis, single-molecule binding traces were tracked, and information of photon counts, and residence time (dwell times) was extracted from kymographs (**Figure 4c**). The dissociation rate constant (k_off_) was determined by fitting a single-exponential decay function (*y* = *A exp*^(−*k t*)^ + *y*0) to the histogram derived from dwell times of 316 single-molecule traces with a bin width of 2 s using Origin (**Figure 4d**). To determine the effect of protein concentration on the k_off_, dwell times from 100, and 106 single-molecule traces were binned to construct histograms at AIM2 concentrations of 1 nM and 5 nM, respectively. Histograms with 2 s bin width were fitted to a single-exponential decay function to determine k_off_ values at the two concentration values (**Figure 4e, f**).

Association rate constants (k_on_) were calculated based on the analysis of 100 single-molecule traces at 1 nM and 109 single-molecule traces at 5 nM in kymographs acquired with a constant time length of 600 s. The unbound times (t_on_) were calculated from the total observation time minus the sum of the residence time of all the traces for each kymograph. The obtained values of t_on_ are summarized in **Supplementary Tables 2 and 3**. The association rates 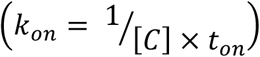were calculated using average t_on_ at 1 nM and 5 nM concentrations. Finally, the dissociation constant (K_D_) was determined from the ratio of the dissociation rate (k_off_) and association rate (k_on_). These results are shown in **Table 1**.

## Acknowledgements

Research reported in this publication was supported by the National Institute of Allergy and Infectious Diseases of the National Institutes of Health under award number R21AI168983 to E.d.A. The content is solely the responsibility of the authors and does not necessarily represent the official views of the National Institutes of Health. E.d.A. and M.S. acknowledge support from the National Science Foundation CREST Center for Cellular and Biomolecular Machines under the award number NSF-HRD-2112675. We are grateful to Dr. Mourad Sadqi for mass spectrometry data acquisition and analysis.

## Data availability

Harbor = https://harbor.lumicks.com

Fiji = https://imagej.net/software/fiji/downloads

OriginLab = https://www.originlab.com

## Supplementary data

A separate file is submitted as supplementary data

## Notes

### Competing Interest Statement

The authors have declared no competing interest.

