## Supplementary Information for "Assembly mechanism of the AIM2 inflammasome sensor revealed by single-molecule analysis"

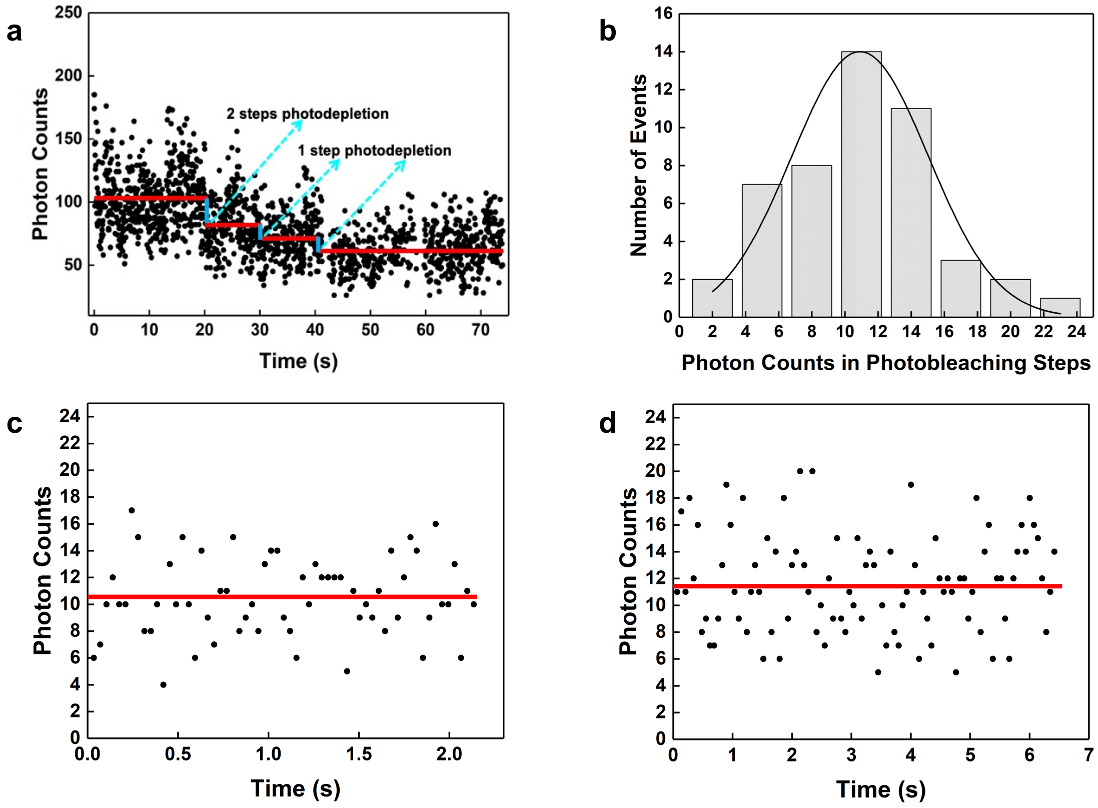


**Supplementary Figure 1. Photodepletion leads to the decrease with time of fluorescence intensity emitted by AIM2 oligomers. a** Example of confocal fluorescence intensity decrease with time due to photodepletion of a single trace extracted from a kymograph. Red lines represent average values of the intensity. The step to the left corresponds to two-fluorophore photodepletion (cyan). The second and third steps correspond to single fluorophore photodepletion (cyan). **b** Distribution of single-step photodepletion events (N = 50) extracted from kymographs for AIM2 oligomers. Fitting to a gaussian function results in an average number of 11 ± 4 photon counts for a single fluorophore. The goodness of the fit is represented by an R-square of 0.95 and RMSE (root mean squared error) of 0.42. **c, d** Examples of two kymographs corresponding to single fluorophores with average photon counts (indicated by horizontal red lines) close to 11 in agreement with the value determined from the distribution in **b**.

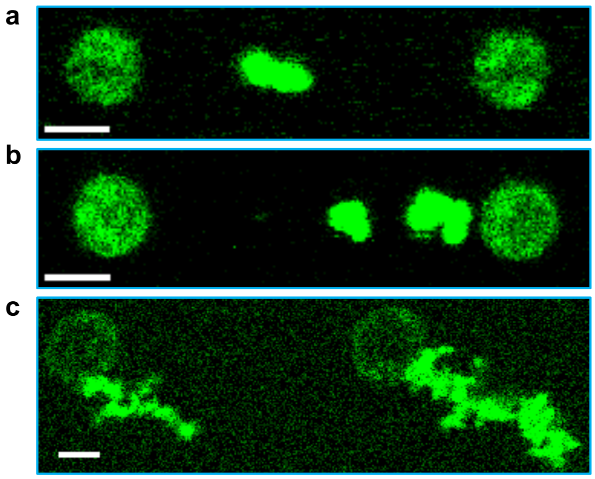


**Supplementary Figure 2. Large AIM2 self-assemblies in the presence and absence of dsDNA. a, b** 2D confocal fluorescence scans showing large AIM2 oligomers bound to the dsDNA (bead diameter and scale bars are 3 µm). **c** 2D confocal fluorescence scans showing large AIM2 oligomers trapped with beads (bead diameter is ~ 4 µm and scale bar is 3 µm).

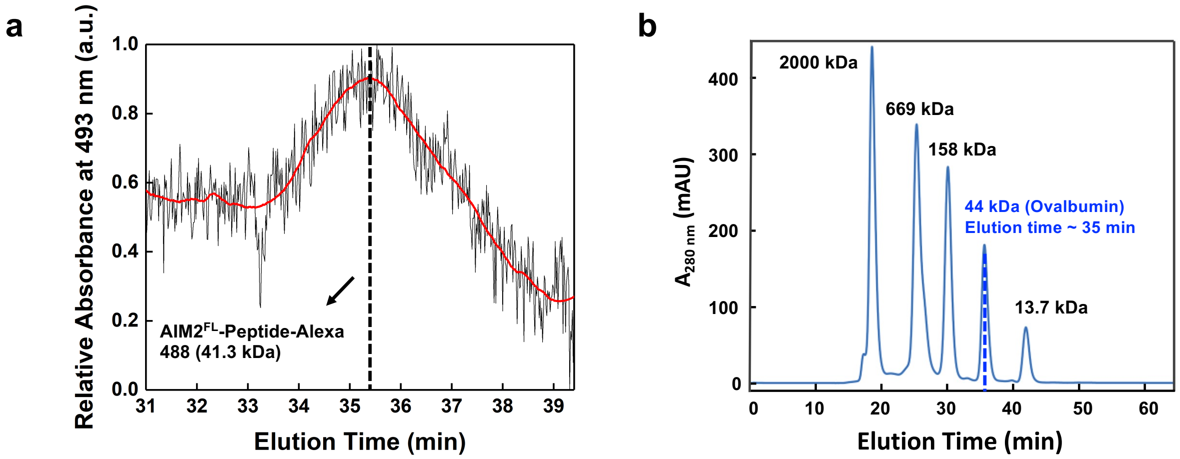


**Supplementary Figure 3. AIM2 is monomeric during purification by size exclusion chromatography. a** Size exclusion chromatogram showing the elution time and absorbance at 493 nm of full-length AIM2 labeled with a peptide carrying Alexa 488 (Matrix: Superdex 200 increase 10/300 GL (GE Healthcare)) after cleavage of 450 nM MBP-AIM2-Peptide-Alexa 488 with TEV protease. **b** Chromatogram reported by GE Healthcare for a mixture of protein standards including ovalbumin with a molecular mass close to AIM2-peptide-Alexa 488. Ovalbumin's elution time is very close to that of AIM2-peptide-Alexa 488.

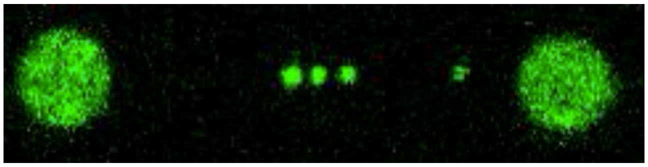

**Supplementary Figure 4. Diffusion of MBP-AIM2 oligomers along the dsDNA.** Movie: Continuous acquisition of 2D scans showing the diffusion of MBP-AIM2 oligomers (250 pM) on dsDNA at 50 mM NaCl.

**Supplementary Table 1**: Permanence times of AIM2 oligomers with different number of protomers on a single dsDNA molecule

| Cluster size (Number of protomers) | Permanence time (s)* |
|---|---|
| 1 | 3.3 |
| 2 | 2.9 |
| 3 | > 680 |
| 4 | > 680 |
| 6 | > 751 |
| 7 | > 627 |
| 8 | > 1516 |
| 9 | > 680 |
| 10 | > 627 |
| 11 | > 619 |
| 14 | > 619 |
| 18 | > 951 |
| 64 | > 1105 |
| 113 | > 751 |
| 205 | > 951 |
| 231 | > 668 |
| 234 | >1516 |
| 309 | > 951 |
| 333 | > 668 |
| 335 | > 643 |
| 339 | > 643 |
| 361 | > 668 |
| 370 | > 951 |
| 514 | > 643 |

* “>” indicates that the oligomer remains attached to the single dsDNA for the total length of the kymograph.

**Supplementary Table 2**: Unbound times ($t_{on}$) of single AIM2 molecules on dsDNA obtained from kymographs acquired at 1 nM protein concentration and 600 s total observation time

| Kymograph | Number of traces analyzed | Unbound time, $t_{on}$ (s) | Average $t_{on}$ (s) |
|---|---|---|---|
| 1 | 26 | 443.2 | 551.0 |
| 2 | 21 | 483.6 | |
| 3 | 8 | 580.7 | |
| 4 | 8 | 519.2 | |
| 5 | 7 | 527.9 | |
| 6 | 5 | 574.8 | |
| 7 | 5 | 528.3 | |
| 8 | 5 | 571.6 | |
| 9 | 4 | 587.4 | |
| 10 | 3 | 584.7 | |
| 11 | 3 | 589.0 | |
| 12 | 3 | 585.4 | |
| 13 | 2 | 587.7 | |

**Supplementary Table 3**: Unbound times ($t_{on}$) of single AIM2 molecules on dsDNA obtained from kymographs acquired at 5 nM protein concentration and 600 s total observation time

| Kymograph | Number of traces analyzed | Unbound time, $t_{on}$ (s) | Average $t_{on}$ (s) |
|---|---|---|---|
| 1 | 14 | 527.6 | 544.3 |
| 2 | 14 | 563.6 | |
| 3 | 13 | 546.6 | |
| 4 | 13 | 528.5 | |
| 5 | 11 | 576.2 | |
| 6 | 11 | 546.5 | |
| 7 | 9 | 524.5 | |
| 8 | 9 | 530.4 | |
| 9 | 8 | 527.1 | |
| 10 | 4 | 581.6 | |
| 11 | 3 | 534.9 | |
